# Reversed upper glycolysis and rapid activation of oxidative pentose phosphate pathway supports the oxidative burst in neutrophils

**DOI:** 10.1101/2020.11.25.396838

**Authors:** Emily C. Britt, Jing Fan

## Abstract

Neutrophils are abundant white blood cells at the frontline of innate immunity. Upon stimulation, neutrophils rapidly activate effector functions such as the oxidative burst and neutrophil extracellular traps (NETs) to eliminate pathogens. However, little is known about how neutrophil metabolism powers these functions. Our metabolomic analysis on primary human neutrophils revealed that neutrophil metabolism is rapidly rewired upon pro-inflammatory activation, with particularly profound changes observed in glycolysis and the pentose phosphate pathway (PPP). We found that the stimulation-induced changes in PPP were specifically coupled with the oxidative burst. The oxidative burst requires a large amount of NADPH to fuel superoxide production via NADPH Oxidase (NOX). Isotopic tracing studies revealed that in order to maximize the NADPH yield from glucose metabolism, neutrophils quickly adopt near complete pentose cycle during the oxidative burst. In this metabolic mode, all glucose is shunted into the oxidative PPP, and the resulting pentose-phosphate is recycled back to glucose-6-phosphate, which then re-enters the oxidative PPP. To enable this recycling, net flux through the upper glycolytic enzyme glucose-6-phosphate isomerase (GPI) is completely reversed. This allows oxidative PPP flux in neutrophils to reach greater than two-fold of the glucose uptake rate, far exceeding other known mammalian cells and tissues. Intriguingly, the adoption of this striking metabolic mode is completely dependent on an increased demand for NADPH associated with the oxidative burst, as inhibition of NOX resets stimulated neutrophils to use glycolysis-dominant glucose metabolism, with oxidative PPP flux accounting for less than 10% of glucose metabolism. Together, these data demonstrated that neutrophils have remarkable metabolic flexibility that is essential to enable the rapid activation of their effector functions.

**Graphic Abstract:** 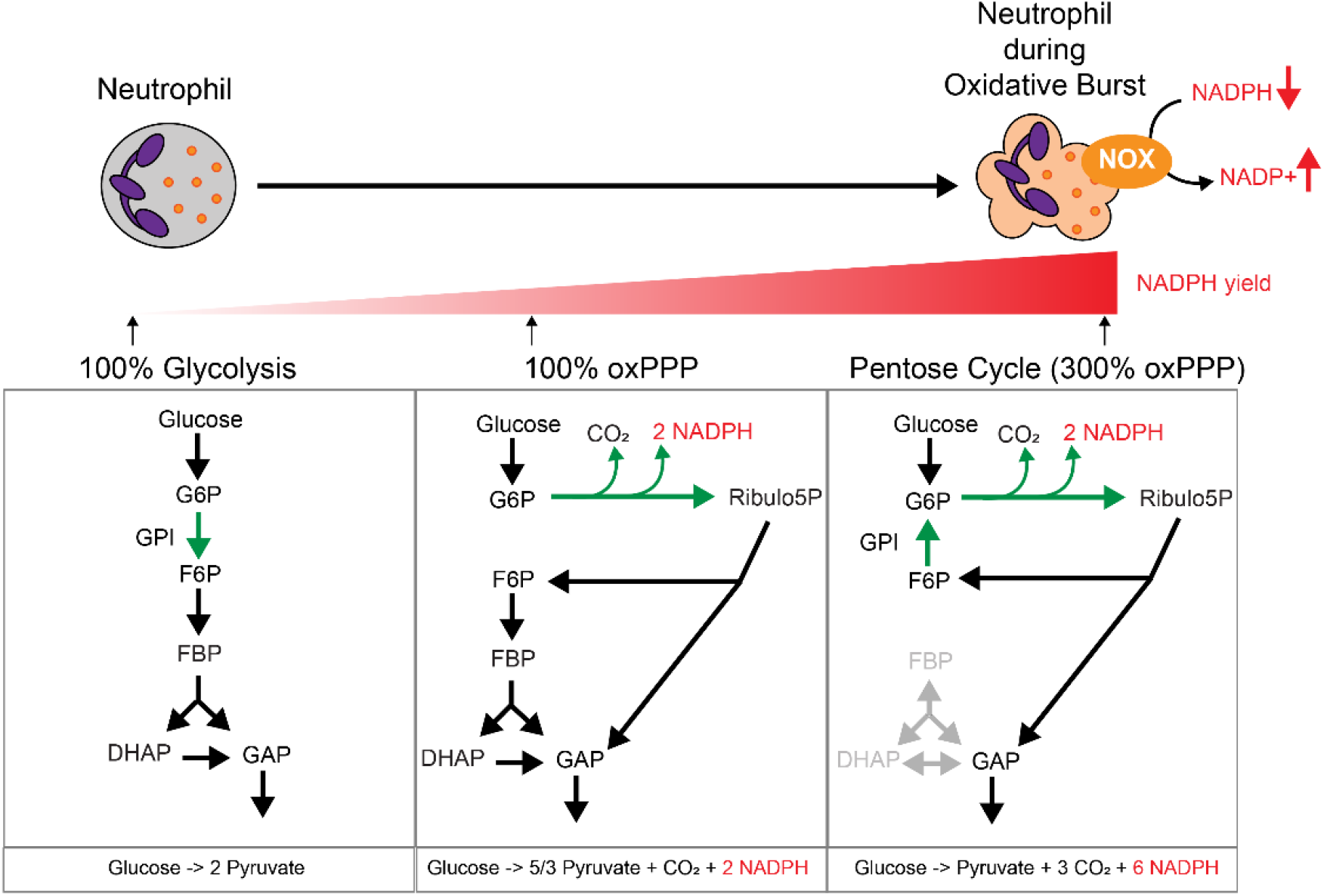

## Introduction

Neutrophils are the most abundant leukocytes in circulation, with a crucial role in innate immunity. During infection, circulating neutrophils migrate towards the infected site upon sensing pathogen-associated molecular patterns (PAMPs) and other molecular signals [1]. They act rapidly to eliminate pathogens with reactive oxygen species (ROS), anti-bacterial peptides and proteases, and neutrophil extracellular traps (NET) [2,3]. The production of ROS is a major component of immune defense in neutrophils, as ROS not only acts directly as a powerful weapon against pathogens, it can also activate granular proteases and induces subsequent NET release [4–6]. The rapid production of ROS upon activation is called an oxidative burst, which is catalyzed by a multi-component enzyme NADPH oxidase (NOX) [7].

Emerging studies have highlighted the vital role of metabolism in both supporting and orchestrating immune functions [8–11]. It has been revealed in several immune cell types, such as T-cells, macrophages and NK cells, that the transition into a specific functional state is often coupled to specific reprogramming of cellular metabolism [12–14]. In contrast, very little is currently known about neutrophil metabolism [15]. In neutrophils, the activation of effector functions is associated with substantial metabolic demands. For instance, the oxidative burst requires a large amount of NADPH to drive the reduction of oxygen. Several recent studies using nutrient perturbation and metabolic inhibitors showed that the oxidative burst and NET release are significantly impacted by glucose metabolism [16–20], demonstrating the importance of metabolism in neutrophil function. However, more comprehensive and in-depth understanding of specific metabolic rewiring associated with neutrophil activation is still lacking. Importantly, as neutrophils are at the first line of immune defense, they have the unique ability to turn on antipathogen functions very quickly. Therefore, a critical question in neutrophil metabolism is to understand how they efficiently utilize limited metabolic resources to support substantial increases in metabolic demand in a very short time.

Here we measured the rapid metabolic changes in primary human neutrophils upon pro-inflammatory activation and identified the specific changes that are directly coupled to the oxidative burst. Using isotopic tracing with a comprehensive set of glucose tracers, we discovered that to support the oxidative burst, neutrophils adopt near complete pentose cycle, a metabolic mode with ultra-high NADPH yield that has not been shown in other mammalian cells. During the oxidative burst, net flux through upper glycolytic enzyme GPI is completely reversed, enabling substantial recycling of pentose carbon and allowing oxidative PPP flux in neutrophils to reach over two-fold the glucose uptake rate. Interestingly, the rapid switch to pentose cycle is entirely on-demand; in the absence of oxidative burst, neutrophils use typical glycolysis as the dominant route for glucose metabolism. This discovery reveals impressive and unique metabolic flexibility in neutrophils that is essential for their function in innate immunity.

## Results

### Primary human neutrophils rapidly reprogram metabolism upon stimulation

To investigate the metabolic alterations associated with neutrophil activation, we isolated primary human neutrophils from the peripheral blood of independent, healthy donors, and stimulated each set of cells with three types of stimulation: (1) Phorbol myristate acetate (PMA), a protein kinase C activator; (2) N-formylmethionyl-leucyl-phenylalanine (fMLP), a bacterial-peptide analog; and (3) the combination of lipopolysaccharide (LPS), a pathogen associated molecular pattern, and interferon-γ (IFNγ), a pro-inflammatory cytokine. These stimulations are commonly used to activate neutrophils and induce pathogen eliminating functions including oxidative burst and NET release [21]. Analysis of metabolite levels in stimulated and unstimulated neutrophils revealed very rapid metabolic rewiring -- just 30 minutes of stimulation was sufficient to induce many significant metabolic alterations that were consistent across donors (Fig.1A). Interesting similarities were observed in the metabolic responses induced by the three stimulations. Particularly, all detected intermediates in the pentose phosphate pathway (PPP) and many intermediates in glycolysis accumulated with all three stimulations. The accumulation in PPP intermediates is the most profound change among measured metabolites, especially upon PMA stimulation (generally over 10-fold), likely because PMA is the most potent and fast-acting stimulation. We therefore focused on the metabolic rewiring included by PMA in subsequent experiments.

**Figure 1.**
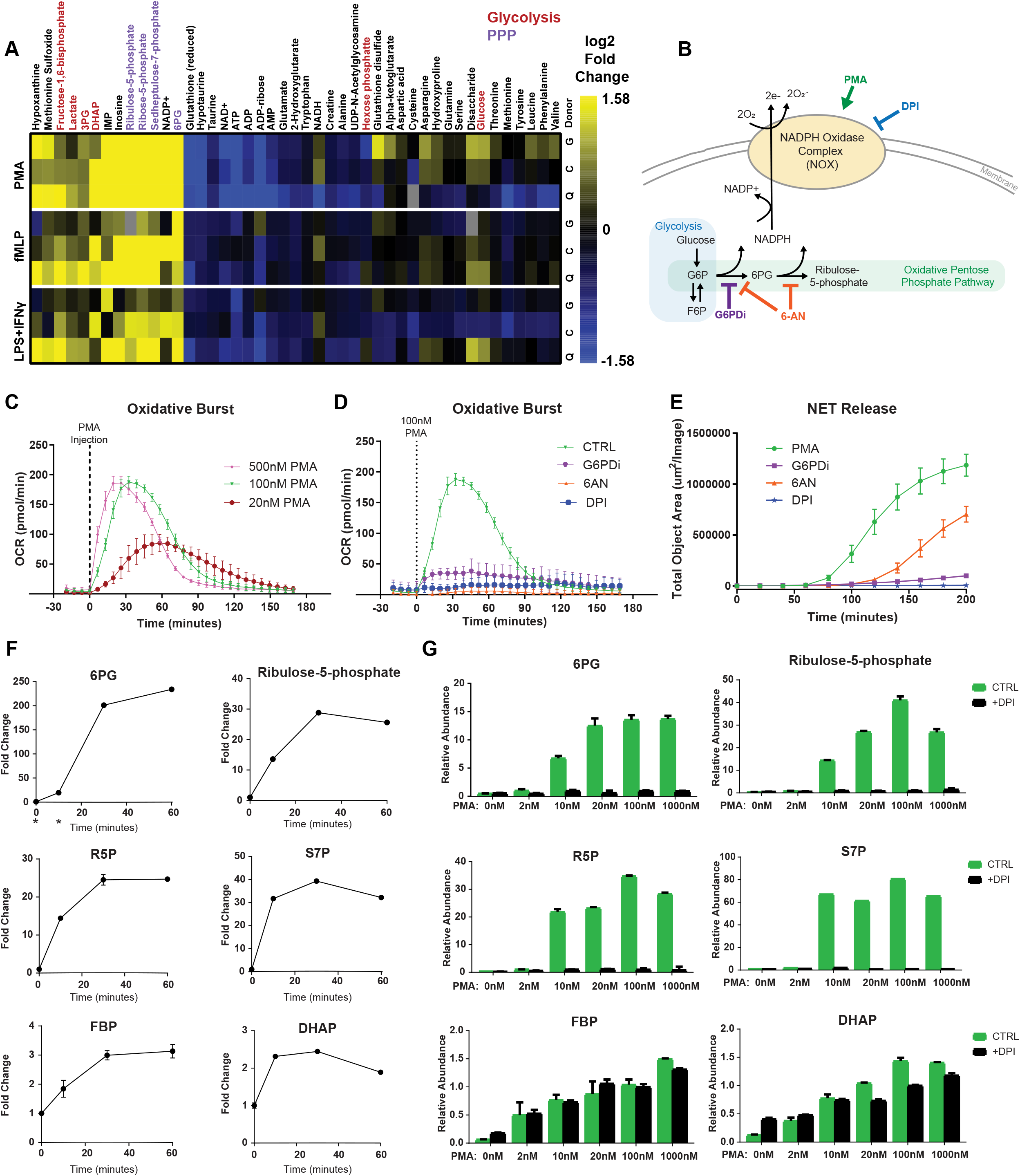
Rapid changes in pentose phosphate pathway is coupled with the oxidative burst. A. Heatmap showing metabolomic changes in primary human neutrophils in response to stimulation of PMA (100nM), fMLP (100nM), or LPS (1μg/ml) and IFNγ (20ng/ml) for 30min. Each row represents results from one donor. Color represents fold change comparing to the unstimulated neutrophils from the same donor, with saturating color indicates 3-fold change or greater. B. Schematic showing the oxidative burst in neutrophils. NADPH oxidase (NOX), which is stimulated by PMA and inhibited by DPI, produces superoxide using NADPH as the electron donor. Oxidative PPP produces NADPH and can by inhibited by G6PDi or 6-AN. C. The oxidative burst induced by varying doses of PMA. Mean ± SD N=6. D. The oxidative burst is inhibited by the treatment of G6PDi (50μM), 6-AN (5mM), or DPI (10μM). Mean ± SD N=6. E. NET release induced by PMA stimulation (100nM added at time 0) is suppressed by the treatment of G6PDi (50μM), 6-AN (5mM), or DPI (10μM). Mean ± SD N=6. F. Changes in the abundances of PPP and glycolytic intermediates over time after stimulation with 100nM PMA. Relative level is normalized to unstimulated neutrophils. * indicates the level in marked condition is below reliable quantitation. G. Relative abundances of specific metabolites after 30min stimulation with varying doses of PMA, with or without the treatment of 10μM DPI.

### Rewiring of the pentose phosphate pathway is coupled to the oxidative burst

As PPP was the most dramatically altered metabolic pathway upon stimulation, we tested how PPP activity affects the activation of major neutrophil effector functions: the oxidative burst and NET release. The oxidative burst can be measured by NOX-dependent increase in oxygen consumption rate (Fig.1B). PMA induced oxidative burst in a dose-dependent manner, with higher dose causing a higher and earlier oxidative burst peak, and the peak plateaued by 100nM PMA (Fig.1C). The oxidative burst has a high demand for NADPH. Inhibiting the oxidative PPP, a NADPH producing pathway, with either 6-aminonicotinamide (6-AN) or glucose-6-phosphate dehydrogenase inhibitor (G6PDi) inhibited the oxidative burst in a dose-dependent manner, almost as completely as directly inhibiting NOX with inhibitor diphenyleneiodonium chloride (DPI) (Fig.1D, Fig S1A,B). NET release starts shortly after oxidative burst ends. NET production was suppressed when NOX was inhibited by DPI (Fig. 1E), suggesting that PMA-induced NET release is largely dependent on NOX produced ROS. Inhibiting oxidative PPP with either G6PDi or 6-AN also partially suppressed NET release in PMA stimulated neutrophils (Fig.1E, Fig S1C). These results are consistent with previous findings [16–18,20,22], indicating that oxidative PPP is required for the oxidative burst, and supports NET release, in activated neutrophils.

We then sought to understand if the observed changes in PPP and glycolysis are coupled to the oxidative burst. As the oxidative burst is a time-dependent process whose magnitude varies depending on the dose of stimulation (Fig.1C), we examined to what extent the metabolite levels change in correlation with the oxidative burst over time and across different dosages of PMA. Upon stimulation of 100nM PMA, significant accumulation of metabolites in PPP and glycolysis occurred as soon as 10 minutes, and generally reached peak level by 30 minutes (Fig.1F), when the oxidative burst reached its peak. With increasing dose of PMA, metabolites in PPP also increased, and reached maximal at 20-100nM PMA, similar to the dose response of oxidative burst (Fig 1G). Treatment of NOX inhibitor DPI completely prevented the accumulation of PPP metabolites at any PMA dose. In contrast, glycolytic intermediates, such as fructose-1,6-bisphosphate (FBP) and dihydroxyacetone phosphate (DHAP), increased with increasing PMA dose, but were largely unaffected by NOX inhibition (Fig 1G). These data strongly suggest that the stimulation-induced accumulation in PPP, but not glycolysis, metabolites are directly coupled to the oxidative burst. This profound accumulation of PPP intermediates is consistent across donors, and is dependent on the activity of oxidative PPP, as inhibiting oxidative PPP with G6PDi significantly dampens the PMA-induced accumulation of PPP intermediates, but not glycolytic intermediates (Fig S2).

In sum, these data illustrated specific coupling between rewiring of PPP and the oxidative burst— the oxidative burst requires oxidative PPP activity, and the rewiring of PPP is dependent on the activation of oxidative burst.

### Neutrophils switch to using oxidative pentose phosphate pathway as the dominant route for glucose metabolism during the oxidative burst

To directly measure changes in metabolic flux through PPP, we fed cells with 1-^13^C-glucose and traced the labeling of downstream metabolites. The oxidative PPP specifically releases the C1 carbon of G6P as CO_2_, while glycolysis retains this carbon. Therefore, if 1-^13^C-glucose is metabolized via glycolysis, three-carbon lower glycolytic metabolites, such as DHAP and lactate, would be 50% unlabeled and 50% 1-labeled, due to the cleavage of fructose-1,6-bisphosphate by aldolase. Alternatively, if 1-^13^C-G6P is metabolized via the oxidative PPP, and the resulting unlabeled pentose-phosphate returns to glycolysis via non-oxidative PPP, it would generate unlabeled lower glycolytic metabolites (Fig. 2A)[23]. In order to understand the rewiring of glucose metabolism that is specifically coupled to the oxidative burst, rather than a general result of stimulation, we compared the tracing results in neutrophils under two conditions: in both conditions neutrophils were stimulated with 100nM PMA for 30min, with one group treated with NOX inhibitor DPI, which serves as the no oxidative burst control, and the second group without the treatment of DPI, which corresponds to peak oxidative burst. We found that without oxidative burst (PMA+DPI), DHAP and lactate were close to 50% labeled, indicating glycolysis was the major route for glucose metabolism. However, during oxidative burst (PMA), this labeled fraction dropped sharply to below 5%. Treatment of 50μM G6PDi, which partially inhibits oxidative PPP, partially prevented the loss of labeling (Fig. 2B). To verify if the loss of labeling is specific to the C1 position, we also performed parallel U-^13^C-glucose labeling as a control, and found in all the conditions, uniformly labeled glucose labeled lower glycolytic intermediates substantially (>90%) (Fig. 2C). Together these data show that oxidative burst causes neutrophils to switch from using glycolysis to using oxidative PPP as the dominant route for glucose metabolism.

**Figure 2.**
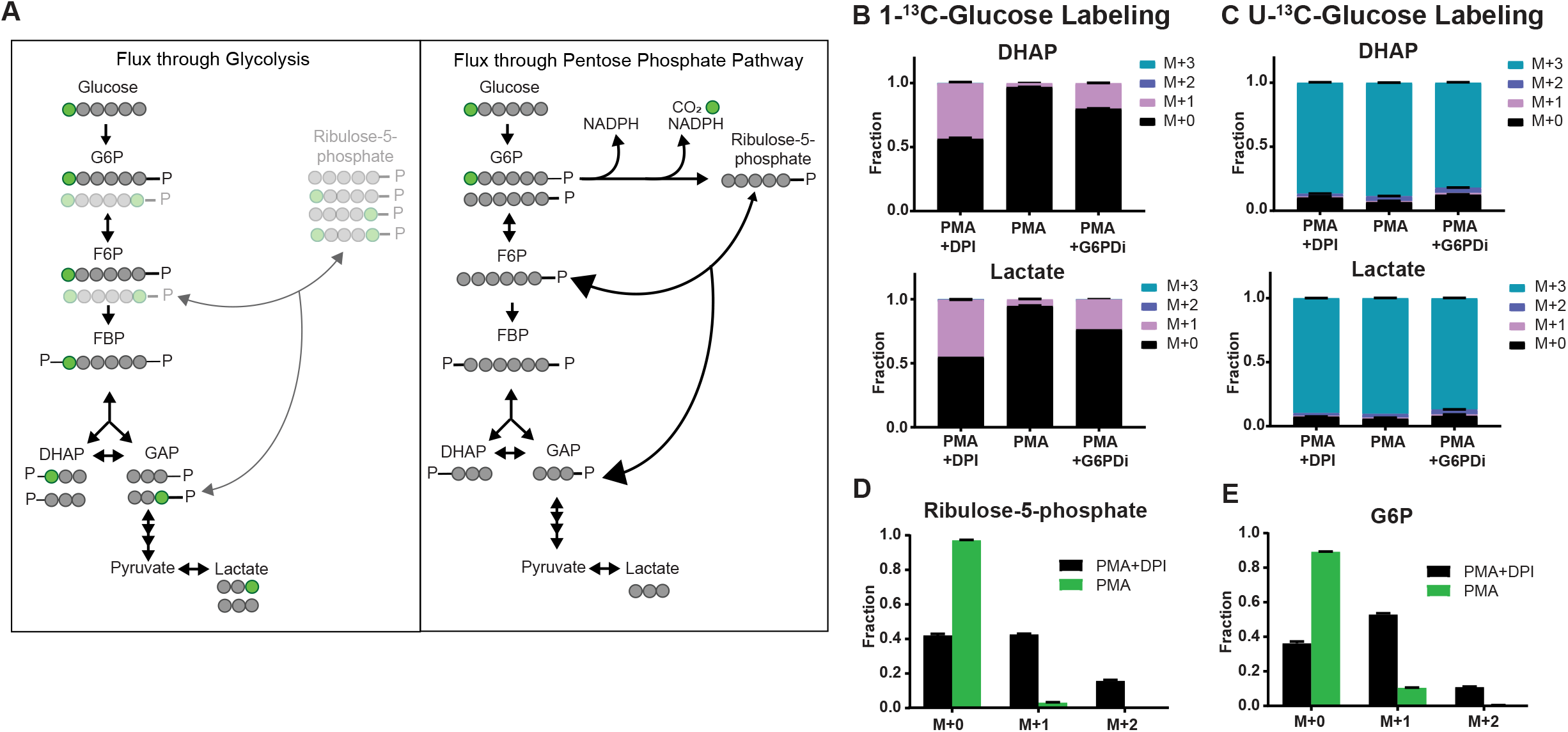
Neutrophils switch to using oxidative PPP as the dominant route for glucose metabolism during the oxidative burst. A. Schematic illustrating major expected isotopomers when 1-^13^C-glucose is fully metabolized via glycolysis (left) or oxidative PPP (right). Some isotopomers generated from reversible activity are shown in grey shade. B, C. Labeling pattern of DHAP and lactate from 1-^13^C-glucose (B) or U-^13^C-glucose (C). D, E. Labeling pattern of ribulose-5-phosphate (D) and G6P (E) from 1-^13^C-glucose in neutrophils with or without oxidative burst.

This labeling study also revealed significant changes in the source of PPP and glycolytic intermediates. Without oxidative burst, pentose-phosphates were slightly above 50% labeled (Fig. 2D, Fig S3A). Since flux through the oxidative PPP produces unlabeled of pentose-phosphates, and flux through the non-oxidative PPP results in mainly 1 or 2-labeled pentose-phosphate, this result shows that in the absence of oxidative burst, both oxidative and non-oxidative PPP contribute significantly to the production of pentose-phosphates. However, during oxidative burst, pentose-phosphates were very close to 100% unlabeled (Fig. 2D, Supp Fig3A), suggesting in this condition oxidative PPP makes the greatest net contribution to pentose production. The overflow of pentose-phosphates from oxidative PPP can be converted by non-oxidative PPP to F6P and GAP, which can be further metabolized via glycolysis. We found that F6P also showed significant loss of labeling during the oxidative burst (Supp Fig 3B), suggesting the non-oxidative PPP contributes significantly to the production of F6P. Very intriguingly, the significant loss of labeling is also observed in G6P (Fig. 2E), indicating that unlabeled F6P from non-oxidative PPP is likely recycled back to G6P via reversed GPI activity.

### Neutrophils recycle carbon via reversed GPI flux to maximize NADPH yield from glucose

The 1-^13^C-glucose tracing revealed that during oxidative burst essentially all G6P is shunted into oxidative PPP instead of glycolysis and pointed to the intriguing possibility of substantial carbon recycling via reversed upper glycolysis. As this recycling is beyond what 1-^13^C-glucose tracing can reliably measure, due to near complete lack of labeling downstream of pentose-phosphates, we further probed the re-distribution of glucose metabolism flux using a 1,2-^13^C-glucose tracer. When this tracer enters the oxidative PPP, the labeled C1 carbon is cleaved and released as ^13^CO_2_, resulting in less labeling enrichment in downstream compounds. If carbon recycling occurs, each glucose molecule can enter the oxidative PPP for more than one round, thus there is a chance to additionally release the C2 carbon as ^13^CO_2_, resulting in even lower labeling enrichment in downstream metabolites. Here we consider three extreme scenarios to analyze the expected average labeling enrichment, i.e. the average number of labeled atoms per molecule, in endproducts pyruvate and lactate (Fig 3A). (i) Full glycolysis: When 100% of the 1,2-^13^C glucose is metabolized via glycolysis with no oxidative PPP activity, all labeled carbon is retained in downstream metabolites. Each 1,2-^13^C glucose is converted to one unlabeled pyruvate (C4-C6) and one 2-labeled pyruvate molecule (C1-C3), thus the average labeling enrichment in pyruvate and lactate would be (0+2)/2=1. (ii) Full non-cyclic oxidative PPP: In this scenario, oxidative PPP flux is 100% of glucose uptake rate, there is no conversion between G6P and F6P. For every three molecules of 1,2-^13^C glucose, three labeled carbons are lost as CO_2_ via the oxidative PPP, resulting in three 1-labeled pentose-phosphates. These 3 pentose phosphates (15 carbons total with 3 labeled) are subsequently converted by non-oxidative PPP to 1 GAP and 2 F6P, which can be further metabolized via glycolysis to 5 pyruvate (still 15 carbons total with 3 labeled). This results in an average labeling of 3/5=0.6. (iii) Full pentose cycle: In this situation, similar to scenario (ii), all G6P is metabolized via the oxidative PPP. However, instead of entering lower glycolysis, the F6P produced from the non-oxidative PPP is all recycled back to G6P via reversed GPI activity. The G6P can then re-enter the oxidative PPP, losing additional labeled carbons as CO_2_. As such, average labeling enrichment in pyruvate or lactate will be lower than that observed in scenario (ii), i.e. <0.6, with the exact number depending on the degree of carbon scrambling in the non-oxidative PPP. In this scenario, each molecule of glucose can go through oxidative PPP three times, with a net production of 3 CO_2_, one pyruvate, and 6 NADPH (Fig 3A). This is the maximal NADPH yield per glucose molecule possible in cells without substantial activity of fructose-1,6-bisphosphatase (FBPase), which is a gluconeogenetic enzyme necessary to fully recycle GAP. We found that the average labeling enrichment in lactate is below 0.3 in stimulated neutrophils during oxidative burst, indicating cells engage in pentose cycle with substantial reversed GPI activity and oxidative PPP flux is greater than 100% of glucose uptake. In contrast, in absence of the oxidative burst, the average labeling enrichment in lactate is very close to 1 (Fig 3B), indicating glycolysis is the major route of glucose metabolism in this condition.

**Figure 3.**
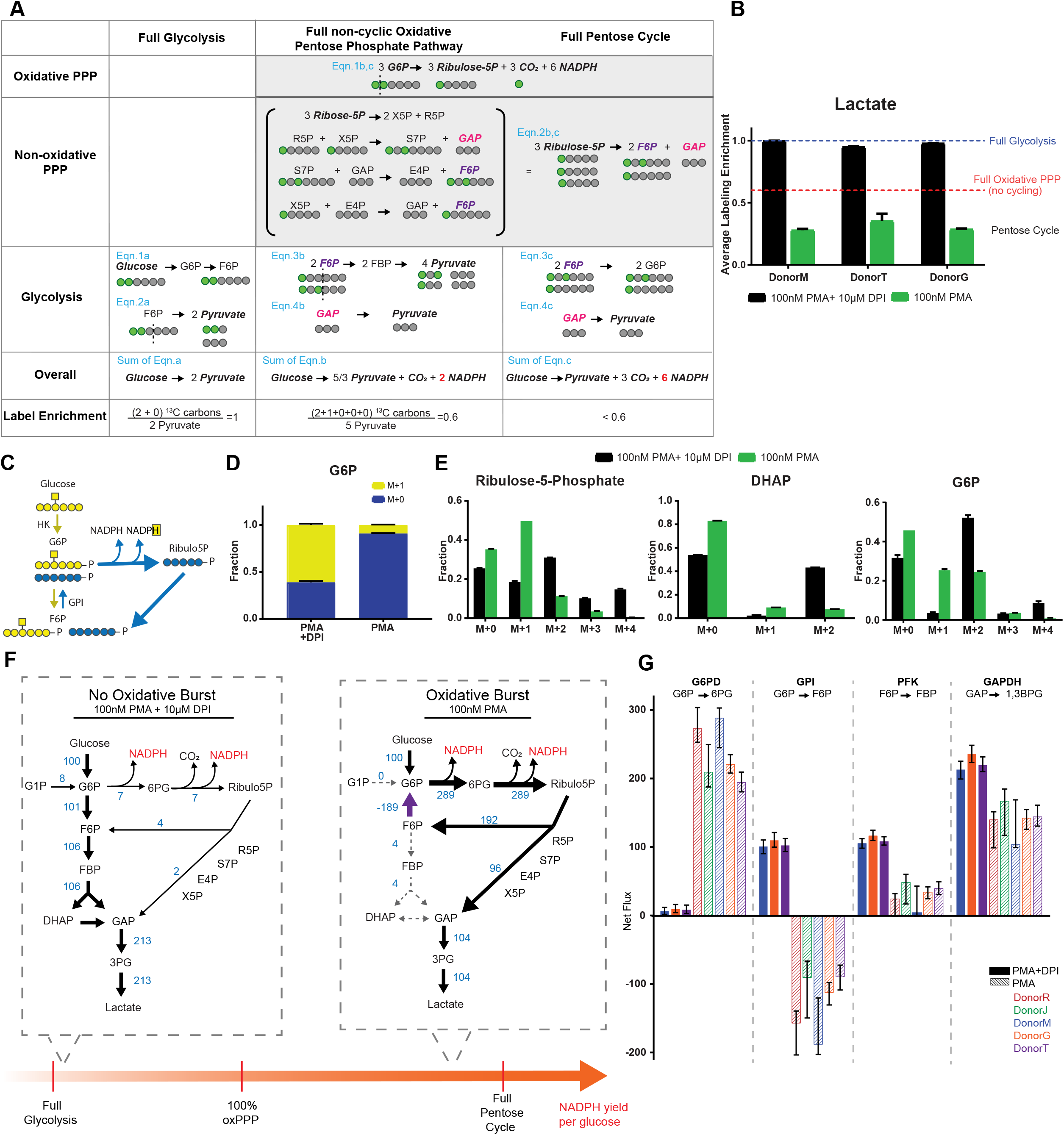
Neutrophils adopt pentose cycle during oxidative burst. A. Schematic illustrating the net reactions, NADPH yield, and expected labeling patterns from 1,2-^13^C-glucose, when cells metabolize glucose 100% via glycolysis, 100% via oxidative PPP, or via pentose cycle with fully reversed GPI net flux (oxidative PPP flux is 300% of total glucose metabolism). B. Average labeling enrichment of lactate from 1,2-^13^C-glucose, measured in neutrophils from 3 donors, with or without oxidative burst. C. Schematic showing hexose kinase (HK) activity generates ^2^H-labeled G6P from 3-^2^H-glucose, whereas pentose cycle generates unlabeled G6P. D. Labeling pattern of G6P from 3-^2^H-glucose. E. Labeling patterns of selected metabolites from 1,2-^13^C-glucose. F. Metabolic flux distributions in neutrophils from one representative donor with or without oxidative burst. Numbers next to the reaction indicate net flux, which are normalized to HK flux as 100. Dotted line indicates the net flux of the reaction is not significantly different from 0 (Zero flux is within confidence interval range). G. Relative net flux of key reactions determined in different donors (indicated by different colors). Solid bars are the fluxes in absence of oxidative burst (30min after stimulation with 100nM PMA, with 10μM DPI treatment) and hatched bars are the fluxes during oxidative burst (30min after stimulation with 100nM PMA). Error bars indicate the confidence intervals from each flux analysis.

To further verify the carbon recycling during oxidative burst, we performed 3-^2^H-glucose labeling. This tracer can be directly converted to ^2^H labeled G6P by hexose kinase (HK). However, when cells engage in pentose cycle, the deuterium at the C3 position is transferred to NADPH when G6P first entered oxidative PPP, therefore, G6P generated by recycling via reversed GPI is unlabeled (Fig 3C). We found that without oxidative burst, the majority of G6P is deuterium labeled, while during oxidative burst, G6P is mostly unlabeled (Fig 3D). This result further supported that during oxidative burst neutrophils adopt substantial pentose cycle.

In addition to average labeling enrichment, we measured the specific labeling patterns of various glycolysis and PPP intermediates from 1,2-^13^C-glucose (Fig 3E, FigS4), which represented the features of glucose metabolism in neutrophils. A few key observations are highlighted below. In pentose-phosphates, we observed a large increase in 1-labeled fraction during oxidative burst, suggesting that oxidative PPP becomes the major route for ribose production. In lower glycolytic compounds like DHAP, the M+1/M+2 ratio increased over 20-fold during oxidative burst, indicating a drastic increase of oxidative PPP activity, which uniquely separates C1 from C2 [24]. In G6P, we saw a drastic increase in the 1-labeled fraction during oxidative burst. This is consistent with substantial carbon recycling. Furthermore, we noticed that without oxidative burst, there is a significant fraction of G6P that is 4-labeled or unlabeled, in addition to the 2-labeled form that directly results from HK activity. This suggests GPI and non-oxidative PPP are highly reversible even at baseline. These interpretations are intriguing while qualitative. We next sought to get a quantitative understanding from these labeling results.

We performed metabolic flux analysis based on measured labeling patterns of eight glycolytic and PPP intermediates using Isotopomer Network Compartmental Analysis (INCA) platform [25]. The simulation yielded a nice fit with the experimental data (Fig S5) and revealed substantial redistribution of glucose metabolism flux coupled to oxidative burst (Fig 3F). To identify features of neutrophil metabolism that are consistent across donors, we performed this analysis in neutrophils from 5 independent donors (in 3 of them we were also able to include DPI treated control). Selected key net fluxes are shown in Fig 3G, and the full results are presented in Table S1. We found that coupled to the activation of oxidative burst, oxidative PPP flux increased from less than 10% to over 200% of glucose uptake rate, and GPI net flux flipped from almost 100% relative to glucose uptake towards F6P to more than 100% reversed (Fig 3F,G). Meanwhile, glycolysis is nearly uncoupled, the net flux from F6P to GAP dropped close to zero, and lower glycolysis (GAP->lactate) became mainly supplied by PPP-derived GAP. This suggests during the oxidative burst neutrophils approach the maximal NADPH yield that can be generated from glucose. Together, this study revealed with high confidence that oxidative burst is coupled to a drastic shift from glycolysis-dominant glucose metabolism to near complete pentose cycle.

### The switch to pentose cycle is rapid and correlated with the magnitude of oxidative burst

We next investigated how the re-distribution of glucose metabolism flux, which is indicated by the changes in average labeling enrichment in lactate from 1,2-^13^C-glucose, occurs over time upon stimulation. We found that as soon as 10 minutes after stimulation, the labeling enrichment in lactate dropped from close to 1, which corresponds to glycolysis-dominant glucose metabolism, to well below 0.6, which indicates substantial pentose cycle (Fig 4A). This shows the switch to pentose cycle occurs very quickly, consistent with data in Fig 1 showing the accumulation of PPP intermediates and the activation of oxidative burst occurs within a short timeframe. The lactate labeling enrichment further decreased to below 0.3 at 30 minutes, when oxidative burst peaks, and raised to slightly below 0.5 at 60 minutes, when the oxidative burst dampened (Fig 4A). This temporal correlation between glucose metabolism rewiring and the oxidative burst suggests that neutrophils can rapidly adjust metabolic strategy and engage PPP to supply NADPH according to the changing demand associated with the oxidative burst.

**Figure 4.**
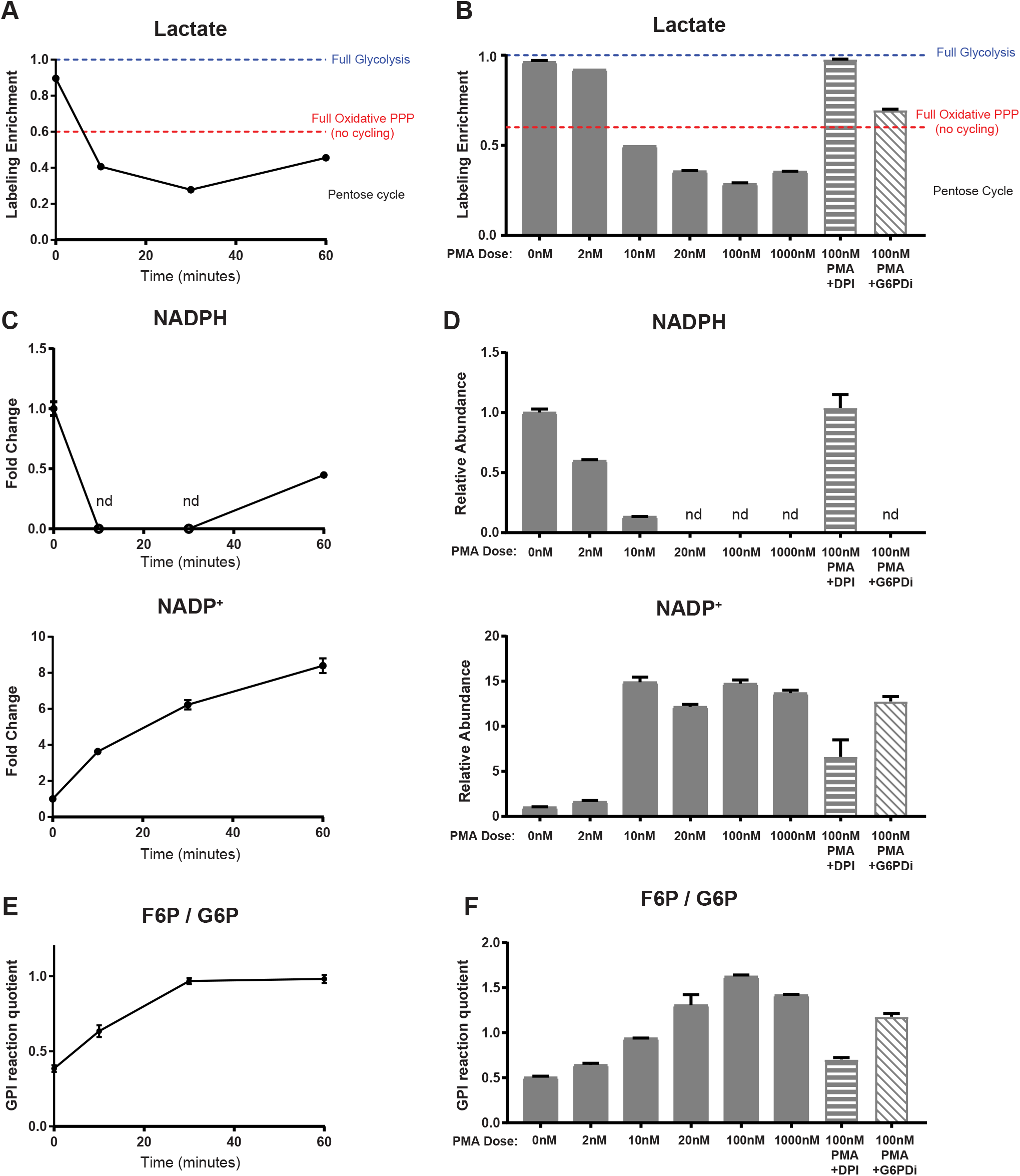
Changes in metabolic state over time and with varying doses of PMA. A, B. Average labeling enrichment of lactate from 1,2-^13^C-glucose changes over time after stimulation with 100nM PMA (A), and with varying doses of PMA at 30min (B). C,D. Relative abundances of NADPH and NADP^+^ change over time after stimulation with 100nM PMA (C), and with varying doses of PMA at 30min (D). E,F. Reaction quotient of GPI, i.e., the ratio between F6P and G6P concentrations, increases over time after simulation with 100nM PMA (E), and changes with increasing dose of PMA or the treatment of 10μM DPI or 50μM G6PDi (F). Mean ± SD. Nd indicates the level is below reliable detection.

We also measured the average labeling enrichment in lactate 30 minutes after stimulation with various doses of PMA. We found labeling enrichment decreased with increasing PMA concentration, and reached minimum when PMA concentration is increased to ~100nM, the dose where peak oxidative burst reaches maximal (Fig 4B, Fig S6). As our data showed neutrophils stimulated with 100nM PMA use near complete pentose cycle, and thus approach the upper limit of NADPH yield from glucose, this likely reflects that maximal metabolic capacity defines the maximal magnitude of oxidative burst. Treating cells with 50μM of G6PDi, which partially inhibited oxidative PPP, partially rescued the lactate labeling enrichment to above 0.6 (Fig 4B), confirming the decrease in labeling indeed reflects the increase of oxidative PPP flux. Eliminating oxidative burst by DPI treatment brings lactate labeling enrichment back to near 1 with any dose of PMA (Fig S6). These results showed that the extent of glucose metabolism rewiring is quantitively coupled with the magnitude of oxidative burst.

### Mechanisms for the switch to pentose cycle

As this study revealed impressive metabolic flexibility in neutrophils, we are interested in the mechanism allowing neutrophils to rapidly switch to pentose cycle on-demand. The labeling results showed this switch can occur within 10 minutes, a timeframe much faster than many mechanisms such as transcriptional regulation typically allows. We therefore reasoned that direct regulation by metabolites plays a major role in driving this metabolic reprogramming.

NADPH and NADP^+^ are classical regulators of oxidative PPP flux. Drop in NADPH/NADP^+^ ratio has been shown to effectively drive the rapid activation of oxidative PPP flux from E.coli to human cells [26–28]. We found that upon PMA stimulation, NADPH dropped to a non-detectable level as soon as 10 minutes, stayed minimal at 30 minutes, and slightly increased by 60 minutes, while NADP^+^ continued to accumulate over time (Fig 4C). Furthermore, the depletion of NADPH and the profound accumulation of NADP^+^ is induced by PMA stimulation in a dose-dependent manner. 20nM PMA was sufficient to cause NADPH to drop below detection and NADP^+^ to increase to maximal level, while inhibiting NOX with DPI fully prevented the depletion of NADPH (Fig 4D). These results showed that NOX activity, which rapidly consumes NADPH and oxidizes it to NADP^+^, dynamically controls NADPH/ NADP^+^ ratio. Upon activation, a drastic decrease in this ratio can act as an indication for NADPH demand and drive a rapid increase in oxidative PPP flux. The strong correlation between NADPH/NADP^+^ ratio and lactate labeling enrichment over time, as well as across PMA doses, supports that the decrease of NADPH/NADP^+^ ratio plays an important role in inducing the switch to pentose cycle in activated neutrophils.

Another key reaction enabling the pentose cycle is reversed GPI activity. This reversibility of GPI is permitted by its thermodynamic status, as GPI reaction has a very a small ΔG^0^ [29]. Our labeling study and metabolic flux analysis revealed significant back and forth GPI gross flux at baseline (Table S1), suggesting GPI rests at a near equilibrium state in neutrophils. Therefore, the net flux direction can be easily flipped by changes in the concentration ratio between F6P and G6P, i.e. the reaction quotient of GPI. During oxidative burst, F6P/ G6P ratio increases in a time-dependent and PMA dose-dependent manner from around 0.5 at baseline to around 1.5 during maximal oxidative burst (Fig 4E,F). This increase is likely due to high oxidative PPP activity that rapidly consumes G6P, and increased non-oxidative PPP flux that produces F6P from the overflowing pentose-phosphates. ROS-driven inhibition of lower glycolytic enzymes may also contribute [30,31]. The theoretical ΔG^0^ of GPI reaction predicts the reaction Kθq is within the range of F6P/G6P in neutrophils. Therefore, the increase in F6P/G6P ratio is sufficient to rapidly reverse the net reaction direction based on thermodynamic control. These results suggest that near-equilibrium upper glycolysis enables the remarkable metabolic flexibility in neutrophils.

## Discussion

Cells can utilize nutrients through different metabolic routes to generate cofactors, energy, and biosynthetic precursors at different proportions according to their needs. Specific to glucose, particular metabolic demands for ATP, NADPH and ribose is best matched with different flux distribution strategies through glycolysis and PPP [32]. Glycolysis produces the most energy and potential energy substrates, at the rate of 2 ATP and 2 pyruvate per glucose. Oxidative PPP produces 2 NADPH and one ribose, while partially oxidizing glucose, reducing its potential to further generate energy. In cells where biosynthetic demand for ribose exceeds the need for NADPH, ribose can be additionally produced from non-oxidative PPP. This occurs in some proliferating cells such as cancer cells [33–35]. When cells have large demand for NADPH but do not need as much ribose, which is the case for many non-proliferating cells like neutrophils, the ribose resulting from oxidative PPP can be shunted back to glycolysis as F6P and GAP via nonoxidative PPP. If the demand for NADPH further exceed the need for other factors, some or all F6P can be recycled back to G6P for additional NADPH production via oxidative PPP. This forms the pentose cycle, which produces 6 NADPH, 1 ATP and 1 pyruvate per molecule of glucose.

The pentose cycle has been proposed in theory as a high NADPH yield glucose metabolism strategy for decades, however, to our knowledge, substantial pentose cycle where oxidative PPP flux exceeds total glucose phosphorylation rate has never been shown in mammalian cells. In typical mammalian cells and tissues, oxidative PPP is usually a minor route for glucose metabolism, with flux less than 10% of glycolysis [36,37]. In conditions involving increased NADPH demands, such as during oxidative stress and fatty acid synthesis, oxidative PPP flux can be higher [26,37–39]. In some of these conditions, there is some carbon recycling at gross flux level [26], i.e., GPI is more reversible, while the net flux is still towards F6P. In neutrophils, without oxidative burst, oxidative PPP flux is 5-10% of glucose uptake, but strikingly, within 30 minutes upon stimulation, it raises to over 2-fold of glucose uptake, far exceeding any known cells and tissues.

Many pathways can potentially contribute to cellular NADPH production. Interestingly, using 3-^2^H-glucose labeling approach [40], we found that without stimulation, active hydride of NADPH barely labels from 3-^2^H-glucose, suggesting oxidative PPP is not the major route for cellular NADPH production at baseline, which is consistent with low oxidative PPP flux measured by ^13^C labeling. The fraction of ^2^H labeled NADPH increased drastically upon PMA stimulation, indicating oxidative PPP becomes the dominant source for NADPH production during oxidative burst (Fig S7). This suggested that neutrophils specifically rely on the upregulation of oxidative PPP, rather than other pathways that also contribute to NADPH production at baseline, to fulfill the large increase of NADPH demand associated with the oxidative burst. It is estimated that in many cell types, due to strong inhibition by NADPH and ATP, and relatively low NADP^+^ availability, oxidative PPP is running at a small fraction of its potential activity [41,42]. This can give cells large reserved metabolic capacity in oxidative PPP to produce NADPH on demand. The metabolite-driven regulation mechanism allows cells to immediately respond to changes in redox status, and the near-equilibrium upper glycolysis and non-oxidative PPP gives neutrophils remarkable flexibility to efficiently use glucose for NADPH production. Together, it allows neutrophils to rapidly power up effector functions and act as the frontline fighters against infection.

The substantial rewiring of PPP is an impressive example of neutrophils’ metabolic flexibility, but it is not the only pathway that changes during neutrophil activation. We have observed other rapid metabolic alterations with each of the stimulations (Fig 1A). Some of these changes, including the changes in glycolytic intermediates, are mainly independent of the oxidative burst (Fig 1G). The accumulation in glycolytic intermediates is likely caused by the translocation of glucose transporter induced by stimulation [16,43]. This can increase glucose uptake rate, which would further increase the supply for oxidative PPP, as well as support other neutrophil functions upon stimulation. Understanding the metabolism in neutrophils is critical for understanding how these important cells work to provide a frontier of innate immunity. This study demonstrated neutrophil metabolism is substantially rewired upon stimulation, and this rewiring is essential for neutrophil functions. Future studies on the reprogramming of a wider range of pathways and its connection with other neutrophil functions will provide a more comprehensive picture of metabolic regulations in neutrophils. This knowledge can also help guide design of metabolic interventions to modulate innate immunity.

## Acknowledgements

We thank Dr. Anna Huttenlocher and David Bennin at University of Wisconsin-Madison for helpful discussions on neutrophil biology. We thank Drs. Tyler Jacobson and Daniel Amador-Noguez at University of Wisconsin-Madison for help on metabolic flux analysis. We thank Dr. Joshua Rabinowitz at Princeton University for generously providing the G6PD inhibitor and helpful discussions. Gretchen Seim and the rest of Fan lab provided valuable feedback on the manuscript. Stephen Halada assisted in phlebotomy. We thank all our blood donors for their participation.

## Methods

### Neutrophil isolation, culture, and stimulation

Human neutrophils were isolated from 8 ml of blood freshly collected from healthy donors, following the protocol approved by the University of Wisconsin Institutional Review Board (protocol number 2019-1031-CP001). Informed consent was obtained from all participants. Neutrophils were purified using the MACSxpress Whole Blood Neutrophil Isolation Kit (Miltenyi Biotec 130-104-434) followed by erythrocyte depletion (Miltenyi Biotec 130-098-196) according to the manufacturer’s instructions. Neutrophils were spun down at 300x g for 5 minutes. Plasma was aspirated off and the cells were resuspended in RPMI-1640 without glutamine (VWR, VWRL0106-0500) supplemented with 5mM L-glutamine (Fisher Scientific, BP379-100) and 0.1% Human Serum Albumin (Lee Biosolutions, 101-15), and kept at 37°C in incubators with 5% CO_2_. Purity of isolated cells was checked by flow cytometry using antibodies against neutrophil surface markers: CD11b (PE) (Biolegend, 301305) and CD15 (AlexFluor700) (Biolegend, 301919), and viability of cells was checked using viability dye Ghost Dye 450 (Tonbo Biosciences, 13-0863). Typically, isolated cells are >90% CD11b+ CD15+ (Fig S8), and > 95% viable. All experiments were performed within 5 hours after blood collection and isolation.

For stimulation, 1.5-3 million neutrophils were aliquoted into 1.5mL Eppendorf tubes and 10ul of 100X stimuli stock solution was added to 1mL cell resuspension. The stimuli was either 1) Phorbol 12-myristate 13-acetate (PMA) (Cayman Chemical, 10008014) 2) N-formylmethionyl-leucyl-phenylalanine (fMLP) (Cayman Chemical, 59880-97-6) or 3) Lipopolysaccharide (LPS) (Sigma-Aldrich L3024) + interferon-γ (IFNγ) (R&D Systems 485-MI). In experiments involving inhibitors, diphenyleneiodonium chloride (DPI) (Sigma-Aldrich, D2926), G6PDi (obtained from Rabinowitz lab at Princeton University [20]), or 6-aminonicotinamide (6-AN) (Cayman Chemical 329-89-5), was added at the same time as stimulation at indicated concentrations.

### Assay for the oxidative burst

The oxidative burst was measured by stimulation-induced oxygen consumption rate using the XF-96e extracellular flux analyzer (Seahorse Bioscience) following protocol developed by Seahorse [44]. To attach neutrophils at the bottom of assay plates (Seahorse Bioscience), neutrophils were plated in culture wells precoated with Cell-Tak (Corning) at 4 × 10^4^ cells per well, and spun at 200x g for 1 minute with minimal acceleration/deceleration, then incubated for 1 h at 37 °C. Assays were performed in regular neutrophil culture media (RPMI-1640 media supplemented with 0.1% human serum albumin). Inhibitors DPI, G6PDi, or 6-AN or vehicle control were added just before starting the assay. After several baseline oxygen consumption rate measurements, PMA was injected through the injection ports to simulate the oxidative burst, and the oxygen consumption rates was continuously monitored throughout the course of oxidative burst.

### NET release assay

NET release was quantified by the increase of extracellular DNA over time following previously developed protocol [45]. Briefly, neutrophils were plated in at 4 × 10^4^ cells per well in 96-well tissue culture plate precoated with Cell-Tak (Corning) and spun at 200x g for 1 minute with minimal acceleration/deceleration. Cytotox Green Reagent (IncuCyte 4633) was added to culture media at 1:4000 to stain extracellular DNA, and images were captured every 20 minutes after stimulation for 5 hours using IncuCyte live cell imager in standard culture condition (37C, 5% CO_2_). Representative images are show in Fig S1C. And the fluorescent area outside of cells, which indicate NET, was quantified by imagine analysis using IncuCyte S3 Basic Analysis software.

### Metabolomics and isotopic tracing

To extract intracellular metabolites, neutrophils were quickly spun at 500x g for 3 minutes at 4C, followed by a quick wash by PBS. Every 2 million of pelleted neutrophils were extracted with 150ul cold liquid chromatography–mass spectrometry (LC–MS) grade acetonitrile:methanol:water 40:40:20 (v:v:v), and the samples were spun at 15,000rpm for 5 minutes at 4C to remove any insoluble debris. The soluble metabolite samples were analyzed using a Thermo Q-Exactive mass spectrometer coupled to a Vanquish Horizon Ultra-High Performance Liquid Chromatograph, using the following two analytical methods. (i) Samples in extraction solvent were directly loaded on to LCMS. The samples were separated on a 2.1 × 150mm XBridge BEH Amide (2.5μm) Column (Waters), using a gradient of solvent A (95% H2O, 5% ACN, 20mM NH_4_AC 20mM NH_4_OH) and solvent B (20% H2O, 80% ACN, 20mM NH_4_AC 20mM NH_4_OH). The gradient was: 0min, 100% B; 3min, 100% B; 3.2min 90% B; 6.2min 90% B; 6.5min 80% B; 10.5min 80% B; 10.7min 70% B; 13.5min 70% B; 13.7min 45% B; 16min 45% B; 16.5min 100% B; 22min 100% B. Flow rate was 0.3ml/ min and column temperature was 30°C. Analytes were measure by MS using full scan. (ii) Samples were dried down under N_2_ flow and resuspended in LC–MS grade water as loading solvent. Metabolites were separated on a 2.1×100mm, 1.7 μM Acquity UPLC BEH C18 Column (Waters) with a gradient of solvent A (97/3 H2O/methanol, 10mM TBA, 9mM acetate, pH 8.2) and solvent B (100% methanol). The gradient was: 0min, 5% B; 2.5min, 5% B; 17min, 95% B; 21min, 95% B; 21.5min, 5% B. Flow rate was 0.2 ml/min. Data were collected with full scan. Settings for the ion source were: 10 aux gas flow rate, 35 sheath gas flow rate, 2 sweep gas flow rate, 3.2 kV spray voltage, 320 °C capillary temperature and 300 °C heater temperature. The identification of metabolites reported here was based on exact m/z and retention times, which were determined with chemical standards. Data were analyzed with Maven.

To determine the molar ratio of F6P / G6P, the concentration of F6P and G6P was quantified using an external calibration curve generated with G6P and F6P standards.

For all experiments involving stable isotope tracers, including 1-^13^C-glucose, 1,2-^13^C-glucose, U-^13^C-glucose, and 3-^2^H-glucose (all from Cambridge Isotope), the isotopically labeled glucose was substituted for unlabeled glucose at the same concentration in RPMI culture media. The natural ^13^C abundance was corrected from the raw data.

### Metabolic Flux analysis

Metabolic flux analysis was performed using the INCA software suite [25]. INCA is implemented in Matlab and simulates isotopic distributions according to the elementary metabolite unit (EMU) framework [46]. We estimated intracellular fluxes by solving a nonlinear least-squares regression problem that minimizes the variance-weighted sum of squared residuals (SSR) between simulated and measured isotopic distributions of intracellular metabolites, under the assumption of pseudo steady state. The network model includes all reactions in glycolysis and PPP, as shown in Table S1. Labeling data from 1,2-^13^C-glucose tracer experiments corrected for naturally occurring heavy isotopes were entered into INCA. Reversible reactions were modeled as a forward and backward reaction. Net fluxes equal forward flux minus backward flux, and exchange flux equals the smaller flux between forward and backward fluxes. Using the optimal solution, we calculated 95% confidence intervals for all estimated fluxes by performing a parameter continuation.

