## Supplemental Figures for "Reversed upper glycolysis and rapid activation of oxidative pentose phosphate pathway supports the oxidative burst in neutrophils"

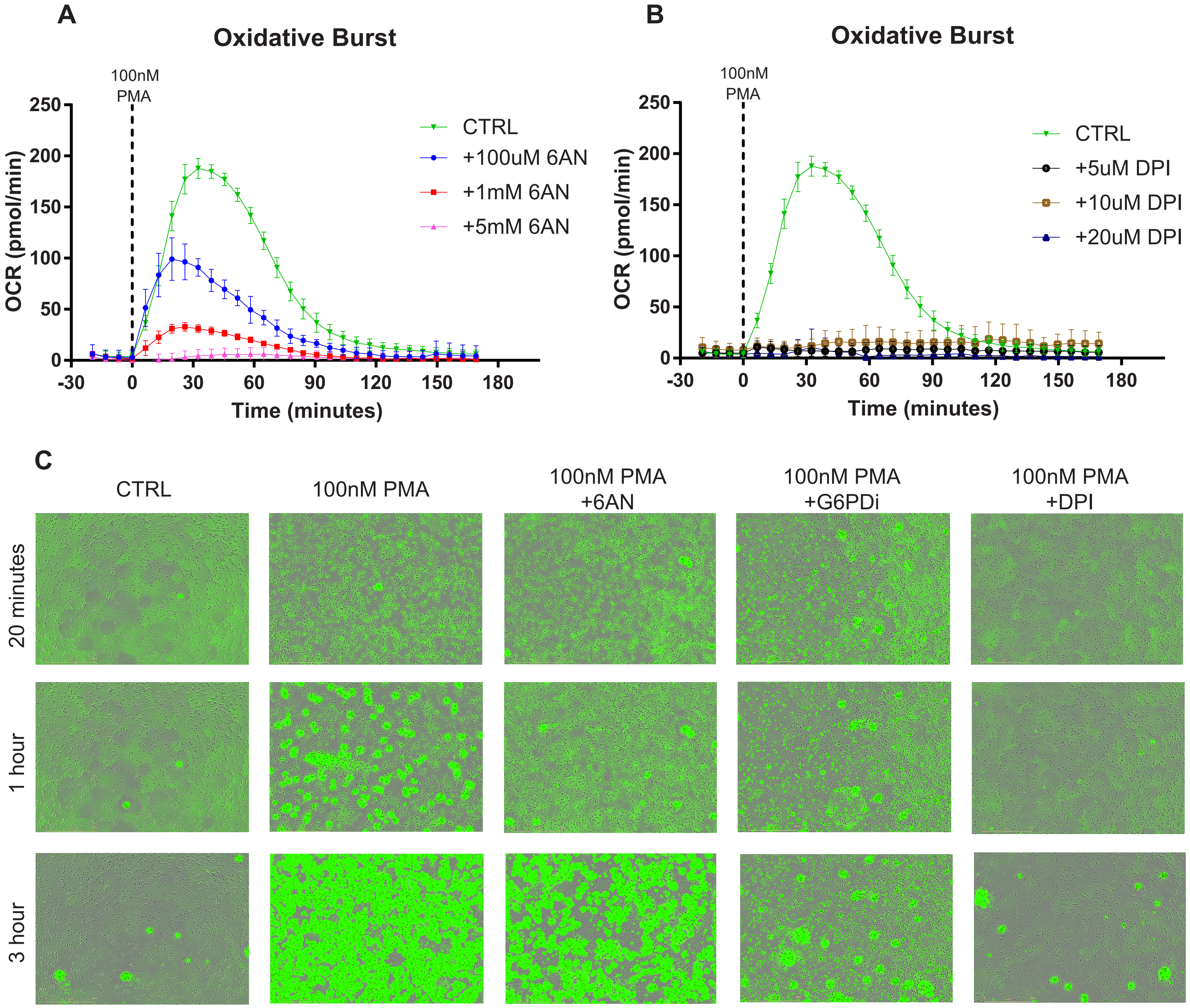


**Figure S1. Oxidative burst and NET release rely on oxidative PPP activity**

A-B. The oxidative burst, which is indicated by PMA induced oxygen consumption rate (OCR), is inhibited by varying concentrations of 6AN (A) or DPI (B). (Mean ± SD, N=6). C. Representative images of NET release by neutrophils stimulated with 100nM PMA, with or without treatment of 6AN (5mM), G6PDi (50µM), or DPI (10µM), at selected time points. NET is indicated by extracellular DNA, which is stained by green fluorescent extracellular DNA dye Incucyte® Cytotox Green Reagent. Images were acquired by IncuCyte live cell imager every 20 minutes, and NET release was quantified and presented in Fig 1E.

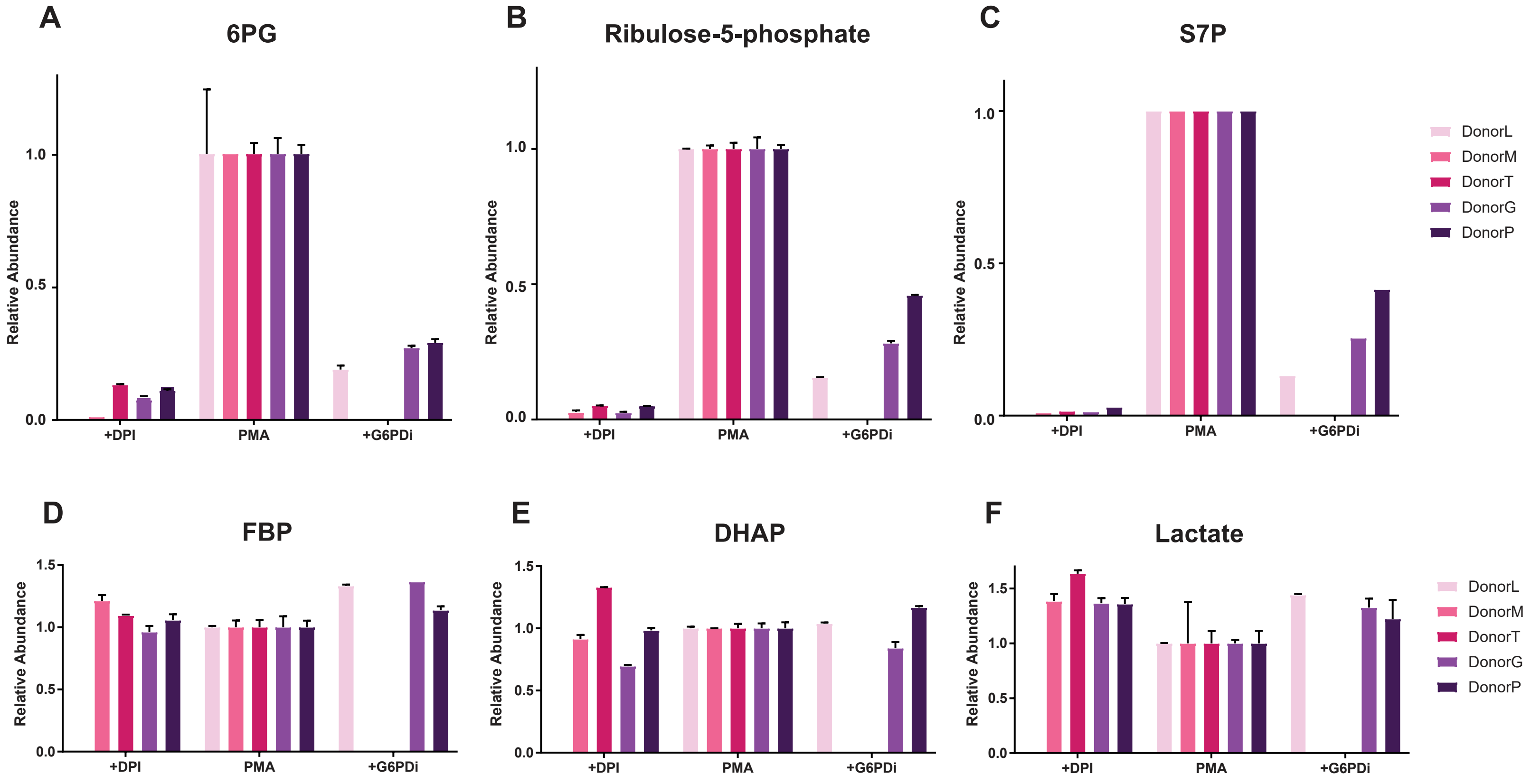


**Figure S2. PMA induced changes in PPP intermediates, but not glycolytic intermediates, are dependent on oxidative burst.**

A-F. Relative levels of PPP intermediates 6PG (A), ribulose-5-phosphate (B), S7P (C), and glycolytic intermediates, FBP (D), DHAP (E), and lactate (F) 30 minutes after 100nM PMA stimulation. Color code indicates different donors. To test if the changes in these metabolites are dependent on oxidative burst, neutrophils isolated from donor M, T, G, P were treated with NOX inhibitor DPI (10μM). To test if the PMA-induced changes in these metabolites are resulted from oxidative PPP activity, some neutrophils from donor L, G, P were treated with G6PDi (50μM). Relative level is normalized to neutrophils stimulated with 100nM PMA without inhibitor from the same donor. Error bars indicate standard deviation between two technical replicates, expect for S7P (C), where only one measurement was made for each donor.

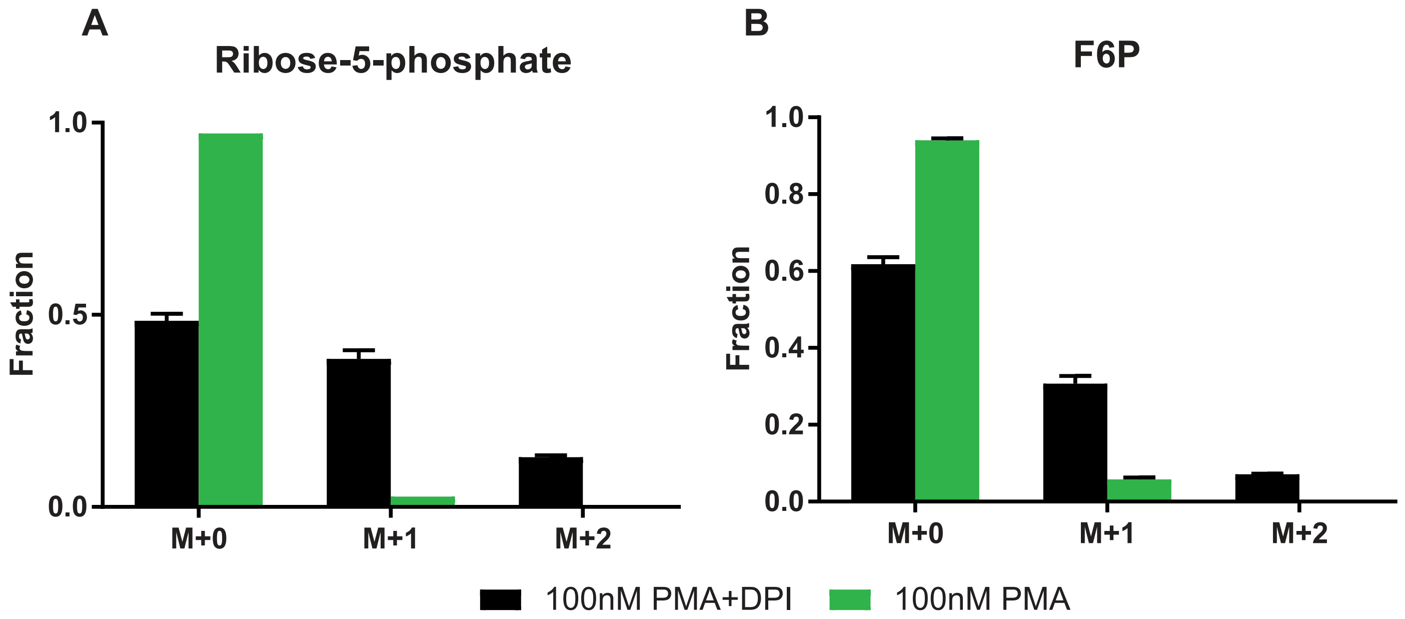


**Figure S3. Labeling pattern of ribose-5-phosphate (A) and fructose 6-phosphate (B) from 1-$^{13}$C glucose.**

Labeling patterns were measured in neutrophils stimulated with PMA (100nM) for 30min, with or without the treatment of NOX inhibitor DPI (10μM). (Mean ± SD). Labeling pattern of selected other metabolites are shown in Fig 2.

*F6P peak may contain some glucose 1-phosphate, as these two compounds are not fully resolved on LCMS.

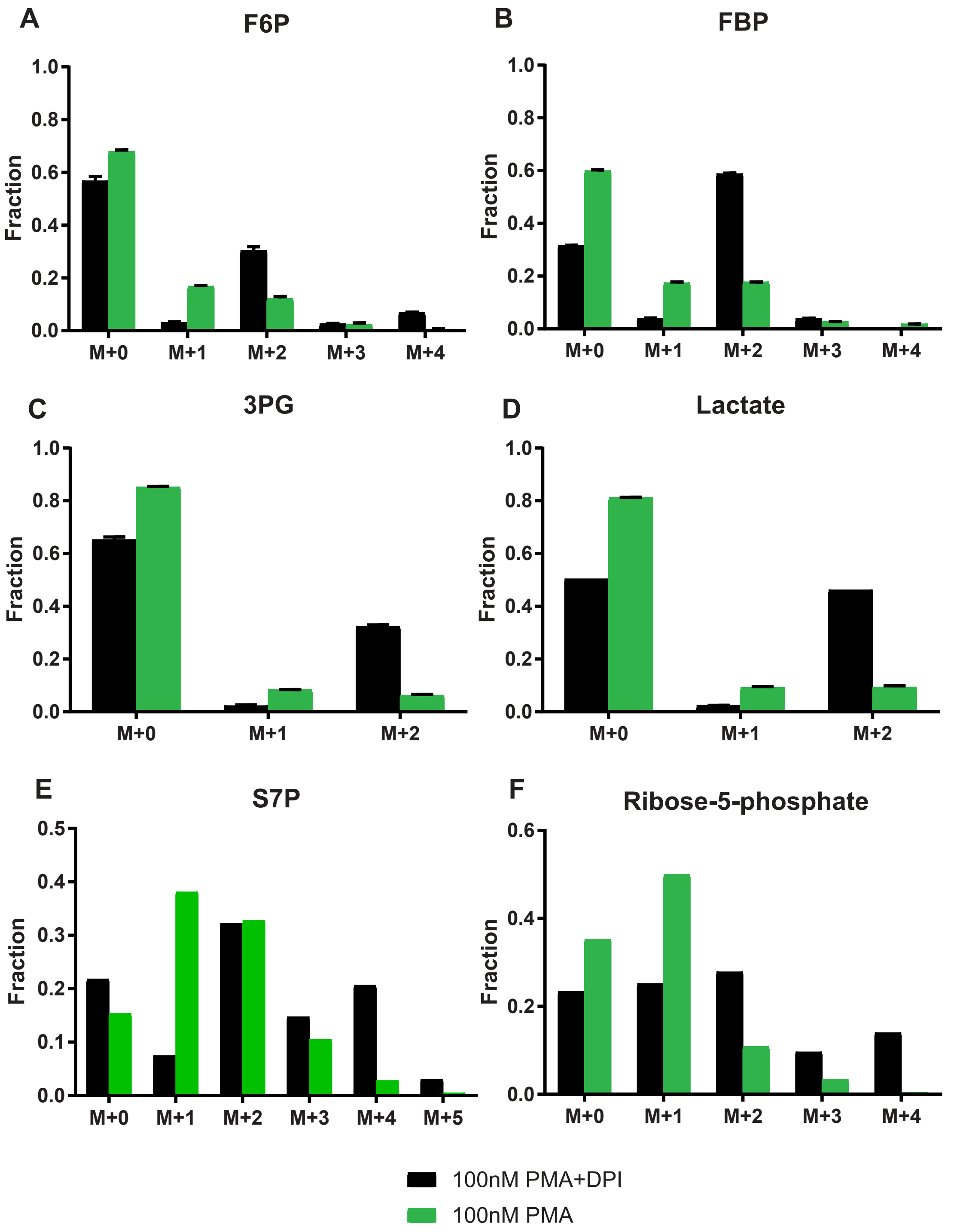


**Figure S4. Labeling pattern of selected PPP and glycolytic intermediates from 1,2-$^{13}C$ glucose.**

Related to Fig 3. Labeling patterns were measured in neutrophils stimulated with PMA (100nM) for 30min, with or without the treatment of NOX inhibitor DPI (10μM). Representative data shown in A-F are from donor G. The same experiment was repeated in neutrophils isolated from other healthy donors (N=5 for PMA stimulated condition, and N=3 for PMA+DPI condition).The results were very similar, and all of these labeling data were used in metabolic flux analysis presented in Fig 3G. F6P (A), FBP (B), 3PG (C), and lactate (D) were measured by LCMS method with a HILIC column. Error bars show standard deviation between technical duplicates. S7P (E), and Ribose-5-phosphate (F) were measured by LCMS method using a C18 column, only one measurement was performed due to limited sample quantity. *F6P peak may contain some glucose 1-phosphate, as these two compounds are not fully resolved on LCMS.

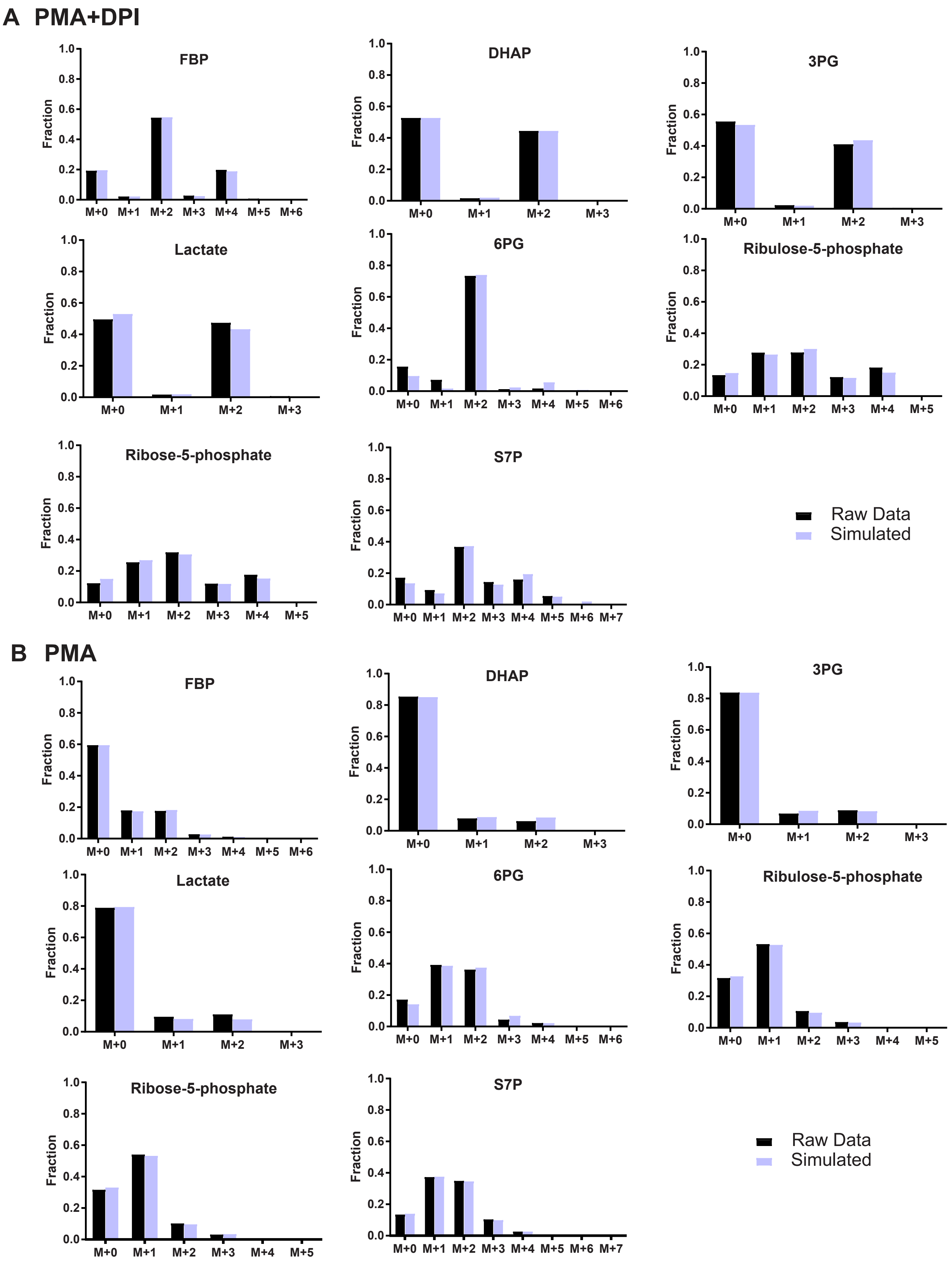


**Figure S5. Comparison between experimentally measured labeling patterns and simulated labeling pattern by the metabolic flux analysis presented in Fig 3F.**

Experimental data (black) and simulation result (blue) are plotted side-by-side in conditions without oxidative burst (A,100nM PMA+ 10μM DPI) or with oxidative burst (B, 100nM PMA). This simulation yielded flux value presented in Fig 3F, with confident interval presented if Fig 3G (Donor M).

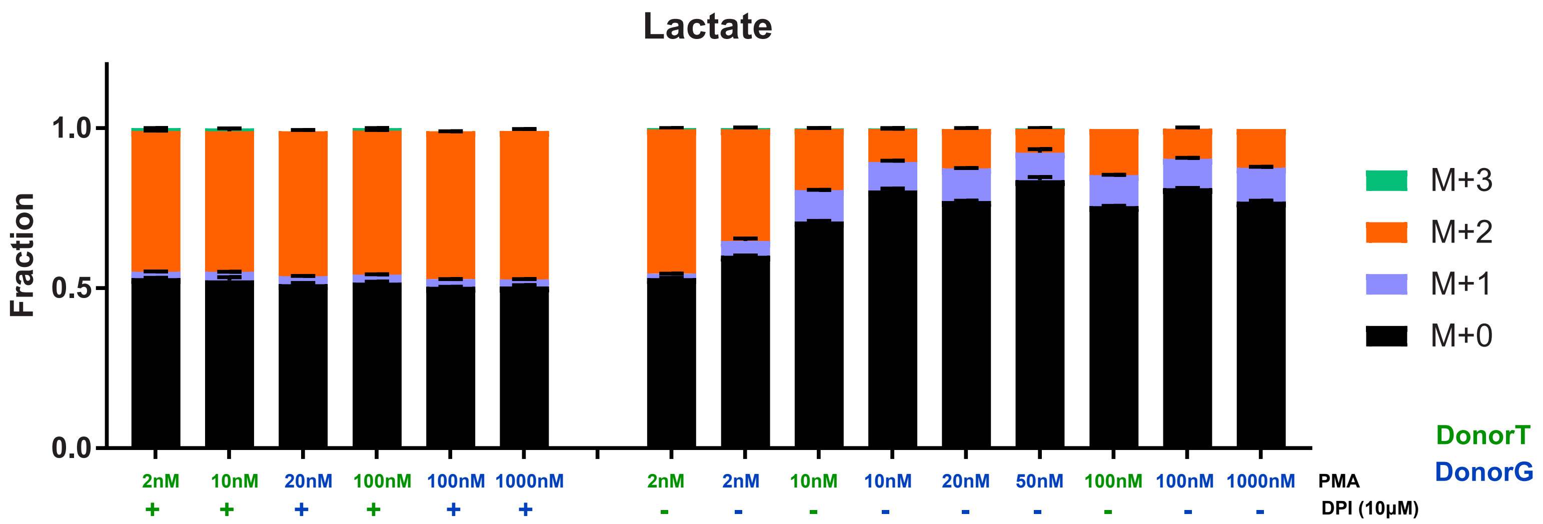


**Figure S6. Lactate labeling pattern from 1,2-$^{13}$C glucose after 30 minutes stimulation with various doses of PMA, with or without the treatment of NOX inhibitor DPI (10μM).**

Due to limited neutrophil yield from each blood draw, the entire span of dose-response curve was covered by neutrophils from two different donors (indicated by color code), with several overlapping conditions (2nM 10nM and 100nM PMA, without DPI) to ensure the trend and main observations are consistent across donors. Mean ± SD between technical duplicates. Related to Fig 4B.

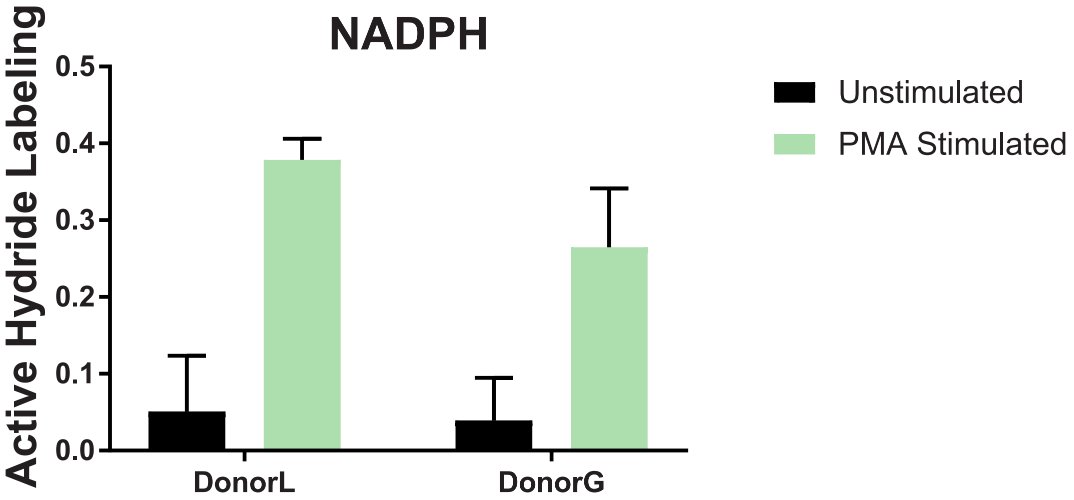


**Figure S7. Active hydride Labeling of NADPH from 3-$^2$H-glucose in unstimulated neutrophils and neutrophils simulated by 100nM PMA for 30min**. Labeling fraction at the active hydride was calculated by comparing the measured isotopic labeling pattern of NADPH to $NADP^+$.

Neutrophil Fraction

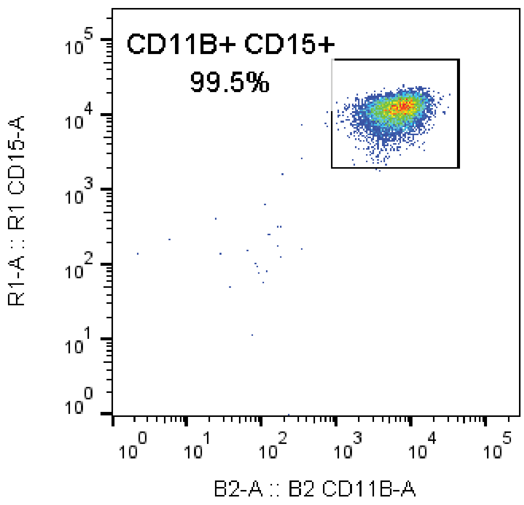


**Figure S8. Quality of isolated primary human neutrophils.** The purity of isolated neutrophils, validated using flow cytometry by surface markers CD15 and CD11B.

| NET FLUX | Donor M 100nM PMA +DPI | | | Donor G 100nM PMA +DPI | | | Donor T 100nM PMA +DPI | | |
|---|---|---|---|---|---|---|---|---|---|
| Eqn | Value | LB | UB | Value | LB | UB | Value | LB | UB |
| Glu.ex -> G6P | 100 | 100 | 100 | 100 | 100 | 100 | 100 | 100 | 100 |
| G6P <-> F6P | 101 | 90 | 110 | 110 | 100 | 121 | 103 | 93 | 112 |
| F6P <-> FBP | 106 | 98 | 112 | 117 | 109 | 124 | 108 | 103 | 115 |
| FBP <-> DHAP + GAP | 106 | 98 | 112 | 117 | 109 | 124 | 108 | 103 | 115 |
| DHAP <-> GAP | 106 | 98 | 112 | 117 | 109 | 124 | 108 | 103 | 115 |
| GAP <-> BPG | 213 | 199 | 225 | 236 | 223 | 248 | 220 | 210 | 231 |
| BPG <-> 3PG | 213 | 199 | 225 | 236 | 223 | 248 | 220 | 210 | 231 |
| 3PG <-> 2PG | 213 | 199 | 225 | 236 | 223 | 248 | 220 | 210 | 231 |
| 2PG <-> PEP | 213 | 199 | 225 | 236 | 223 | 248 | 220 | 210 | 231 |
| PEP <-> Pyr | 213 | 199 | 225 | 236 | 223 | 248 | 220 | 210 | 231 |
| Pyr <-> Lactate | 213 | 199 | 225 | 236 | 223 | 248 | 220 | 210 | 231 |
| Lactate -> Lactate.ex | 213 | 199 | 225 | 236 | 223 | 248 | 220 | 210 | 231 |
| G6P -> 6Pgl | 7 | 2 | 12 | 10 | 4 | 16 | 9 | 3 | 15 |
| 6Pgl <-> 6PG | 7 | 2 | 12 | 10 | 4 | 16 | 9 | 3 | 15 |
| 6PG <-> Ribulo5P + CO2 | 7 | 2 | 12 | 10 | 4 | 16 | 9 | 3 | 15 |
| Ribulo5P <-> R5P | 2 | 1 | 4 | 3 | 1 | 5 | 3 | 1 | 5 |
| Ribulo5P <-> X5P | 4 | 1 | 8 | 6 | 3 | 11 | 6 | 2 | 10 |
| X5P + R5P <-> GAP + S7P | 2 | 1 | 4 | 3 | 1 | 5 | 3 | 1 | 5 |
| X5P + E4P <-> GAP +F6P | 2 | 1 | 4 | 3 | 1 | 5 | 3 | 1 | 5 |
| S7P + GAP <-> E4P + F6P | 2 | 1 | 4 | 3 | 1 | 5 | 3 | 1 | 5 |
| CO2 -> CO2.ex | 7 | 2 | 12 | 10 | 4 | 16 | 9 | 3 | 15 |
| G1P.ex -> G6P | 8 | 1 | 13 | 20 | 14 | 26 | 11 | 7 | 17 |

| EXCHANGE FLUX | Donor M 100nM PMA +DPI | | | Donor G 100nM PMA +DPI | | | Donor T 100nM PMA +DPI | | |
|---|---|---|---|---|---|---|---|---|---|
| Eqn | Value | LB | UB | Value | LB | UB | Value | LB | UB |
| G6P <-> F6P | 45 | 0 | 622 | 159 | 53 | 461 | 162 | 47 | 32780 |
| F6P <-> FBP | 3739 | 118 | Inf | 9999900 | 425 | Inf | 661 | 137 | Inf |
| FBP <-> DHAP + GAP | 186 | 101 | 694 | 345 | 163 | 1048 | 1064 | 459 | 105720 |
| DHAP <-> GAP | 6372 | 1066 | 2206300 | 1025 | 512 | 5921 | 7718 | 1098 | Inf |
| GAP <-> BPG | 50 | 0 | Inf | 1298 | 0 | 1311 | 0 | 0 | Inf |
| BPG <-> 3PG | 498 | 0 | 512 | 479 | 0 | Inf | 6667 | 0 | 6677 |
| 3PG <-> 2PG | 6383 | 0 | 6397 | 2168 | 0 | 2181 | 4562 | 0 | 4572 |
| 2PG <-> PEP | 2032 | 0 | 2046 | 527 | 0 | Inf | 752 | 0 | Inf |
| PEP <-> Pyr | 87 | 0 | Inf | 8956 | 0 | 8969 | 7362 | 0 | 7372 |
| Pyr <-> Lactate | 7859 | 0 | 7873 | 3284 | 0 | 3297 | 9897 | 0 | 9907 |
| 6Pgl <-> 6PG | 9994 | 0 | Inf | 7569 | 0 | 7575 | 5589 | 0 | Inf |
| 6PG <-> Ribulo5P + CO2 | 1 | 0 | 2 | 1 | 0 | 4 | 1 | 0 | 3 |
| Ribulo5P <-> R5P | 1867700 | 0 | Inf | 1051900 | 0 | 1051900 | 0 | 0 | 2247700 |
| Ribulo5P <-> X5P | 10000000 | 10 | Inf | 33 | 8 | Inf | 37 | 9 | 1020 |
| X5P + R5P <-> GAP + S7P | 0 | 0 | 198100 | 0 | 0 | 4620 | 0 | 0 | Inf |
| X5P + E4P <-> GAP +F6P | 30 | 1 | 83 | 2096900 | 4 | 2096900 | 141730 | 15 | 141730 |
| S7P + GAP <-> E4P + F6P | 3 | 1 | 14 | 4 | 1 | 24 | 18 | 1 | 3381700 |

**Table S1. Metabolic flux analysis result. LB, lower bound. UB, upper bound.**

| | Donor R | | | Donor J | | | Donor M | | | Donor G | | | Donor T | | |
|---|---|---|---|---|---|---|---|---|---|---|---|---|---|---|---|
| **NET FLUX** | 100nM PMA | | | 100nM PMA | | | 100nM PMA | | | 100nM PMA | | | 100nM PMA | | |
| Eqn | Value | LB | UB | Value | LB | UB | Value | LB | UB | Value | LB | UB | Value | LB | UB |
| Glu.ex -> G6P | 100 | 100 | 100 | 100 | 100 | 100 | 100 | 100 | 100 | 100 | 100 | 100 | 100 | 100 | 100 |
| G6P <-> F6P | -157 | -204 | -140 | -91 | -150 | -67 | -189 | -203 | -120 | -113 | -131 | -98 | -90 | -109 | -73 |
| F6P <-> FBP | 25 | -1 | 32 | 49 | 17 | 60 | 4 | -1 | 43 | 34 | 24 | 42 | 40 | 30 | 49 |
| FBP <-> DHAP + GAP | 25 | -1 | 32 | 49 | 17 | 60 | 4 | -1 | 43 | 34 | 24 | 42 | 40 | 30 | 49 |
| DHAP <-> GAP | 25 | -1 | 32 | 49 | 17 | 60 | 4 | -1 | 43 | 34 | 24 | 42 | 40 | 30 | 49 |
| GAP <-> BPG | 140 | 99 | 151 | 167 | 117 | 185 | 104 | 99 | 169 | 143 | 126 | 155 | 144 | 130 | 161 |
| BPG <-> 3PG | 140 | 99 | 151 | 167 | 117 | 185 | 104 | 99 | 169 | 143 | 126 | 155 | 144 | 130 | 161 |
| 3PG <-> 2PG | 140 | 99 | 151 | 167 | 117 | 185 | 104 | 99 | 169 | 143 | 126 | 155 | 144 | 130 | 161 |
| 2PG <-> PEP | 140 | 99 | 151 | 167 | 117 | 185 | 104 | 99 | 169 | 143 | 126 | 155 | 144 | 130 | 161 |
| PEP <-> Pyr | 140 | 99 | 151 | 167 | 117 | 185 | 104 | 99 | 169 | 143 | 126 | 155 | 144 | 130 | 161 |
| Pyr <-> Lactate | 140 | 99 | 151 | 167 | 117 | 185 | 104 | 99 | 169 | 143 | 126 | 155 | 144 | 130 | 161 |
| Lactate -> Lactate.ex | 140 | 99 | 151 | 167 | 117 | 185 | 104 | 99 | 169 | 143 | 126 | 155 | 144 | 130 | 161 |
| G6P -> 6Pgl | 273 | 253 | 304 | 210 | 188 | 250 | 289 | 245 | 303 | 221 | 208 | 235 | 195 | 180 | 209 |
| 6Pgl <-> 6PG | 273 | 253 | 304 | 210 | 188 | 250 | 289 | 245 | 303 | 221 | 208 | 235 | 195 | 180 | 209 |
| 6PG <-> Ribulo5P + CO2 | 273 | 253 | 304 | 210 | 188 | 250 | 289 | 245 | 303 | 221 | 208 | 235 | 195 | 180 | 209 |
| Ribulo5P <-> R5P | 91 | 84 | 101 | 70 | 63 | 83 | 96 | 82 | 101 | 74 | 69 | 78 | 65 | 60 | 70 |
| Ribulo5P <-> X5P | 182 | 169 | 202 | 140 | 125 | 166 | 192 | 163 | 202 | 147 | 139 | 156 | 130 | 120 | 140 |
| X5P + R5P <-> GAP + S7P | 91 | 84 | 101 | 70 | 63 | 83 | 96 | 82 | 101 | 74 | 69 | 78 | 65 | 60 | 70 |
| X5P + E4P <-> GAP +F6P | 91 | 84 | 101 | 70 | 63 | 83 | 96 | 82 | 101 | 74 | 69 | 78 | 65 | 60 | 70 |
| S7P + GAP <-> E4P + F6P | 91 | 84 | 101 | 70 | 63 | 83 | 96 | 82 | 101 | 74 | 69 | 78 | 65 | 60 | 70 |
| CO2 -> CO2.ex | 273 | 253 | 304 | 210 | 188 | 250 | 289 | 245 | 303 | 221 | 208 | 235 | 195 | 180 | 209 |
| G1P.ex -> G6P | 16 | 0 | 21 | 19 | 0 | 25 | 0 | 0 | 26 | 8 | 1 | 17 | 5 | 0 | 12 |

| **EXCHANGE** | Donor R | | | Donor J | | | Donor M | | | Donor G | | | Donor T | | |
|---|---|---|---|---|---|---|---|---|---|---|---|---|---|---|---|
| **FLUX** | 100nM PMA | | | 100nM PMA | | | 100nM PMA | | | 100nM PMA | | | 100nM PMA | | |
| Eqn | Value | LB | UB | Value | LB | UB | Value | LB | UB | Value | LB | UB | Value | LB | UB |
| G6P <-> F6P | 148 | 57 | 410 | 421 | 198 | 1893 | 356 | 133 | 1352 | 47 | 0 | 108 | 110 | 54 | 227 |
| F6P <-> FBP | 0 | 0 | 38 | 0 | 0 | 34 | 33 | 0 | 42 | 0 | 0 | 10 | 0 | 0 | 9 |
| FBP <-> DHAP + GAP | 55 | 30 | 112 | 199 | 102 | 653 | 162 | 84 | 595 | 240 | 100 | 5865 | 251 | 109 | 2278 |
| DHAP <-> GAP | 10000000 | 245 | Inf | 1036500 | 614 | 1036500 | 10000000 | 540 | Inf | 10000000 | 554 | Inf | 718000 | 581 | 718010 |
| GAP <-> BPG | 0 | 0 | Inf | 0 | 0 | Inf | 8 | 0 | Inf | 0 | 0 | Inf | 0 | 0 | 1064400 |
| BPG <-> 3PG | 3594 | 0 | Inf | 7304 | 0 | 7356 | 1764 | 0 | 1769 | 131 | 0 | Inf | 429 | 0 | 4229800 |
| 3PG <-> 2PG | 416 | 0 | Inf | 249 | 0 | 300 | 9963 | 0 | 9968 | 1152 | 0 | Inf | 4114 | 0 | Inf |
| 2PG <-> PEP | 471 | 0 | Inf | 64 | 0 | Inf | 1279 | 0 | 1284 | 2367 | 0 | Inf | 1987 | 0 | Inf |
| PEP <-> Pyr | 1310 | 0 | Inf | 0 | 0 | Inf | 24 | 0 | Inf | 125 | 0 | Inf | 5174 | 0 | Inf |
| Pyr <-> Lactate | 244 | 0 | Inf | 1161 | 0 | 1212 | 6187 | 0 | 6192 | 6298 | 0 | Inf | 6465 | 0 | Inf |
| 6Pgl <-> 6PG | 4672 | 0 | 4692 | 9856 | 0 | 9880 | 378 | 0 | 1073900 | 5852 | 0 | Inf | 1407 | 0 | Inf |
| 6PG <-> Ribulo5P + CO2 | 39 | 0 | 153 | 0 | 0 | 63 | 1309 | 415 | Inf | 0 | 0 | 12 | 0 | 0 | 13 |
| Ribulo5P <-> R5P | 27287 | 0 | 27294 | 8066 | 0 | 8074 | 9999900 | 0 | Inf | 9999900 | 929 | Inf | 776730 | 535 | 776740 |
| Ribulo5P <-> X5P | 403 | 0 | Inf | 100730 | 0 | 100750 | 1 | 0 | Inf | 0 | 0 | 284630 | 23 | 0 | Inf |
| X5P + R5P <-> GAP + S7P | 0 | 0 | Inf | 0 | 0 | Inf | 0 | 0 | Inf | 1853600 | 0 | Inf | 9885 | 0 | Inf |
| X5P + E4P <-> GAP +F6P | 210 | 0 | Inf | 71 | 0 | Inf | 0 | 0 | Inf | 0 | 0 | Inf | 3606 | 0 | 555660 |
| S7P + GAP <-> E4P + F6P | 0 | 0 | 248 | 18 | 0 | 1537 | 35 | 0 | Inf | 9999900 | 493 | Inf | 577890 | 513 | Inf |

**Table S1 cont.** Metabolic flux analysis result. LB, lower bound. UB, upper bound.
